# Power and sample size calculations for fMRI studies based on the prevalence of active peaks

**DOI:** 10.1101/049429

**Authors:** Joke Durnez, Jasper Degryse, Beatrijs Moerkerke, Ruth Seurinck, Vanessa Sochat, Russell A. Poldrack, Thomas E. Nichols

## Abstract

**Highlights:**

- The manuscript presents a method to calculate sample sizes for fMRI experiments
- The power analysis is based on the estimation of the mixture distribution of null and active peaks
- The methodology is validated with simulated and real data.

**Abstract:** Mounting evidence over the last few years suggest that published neuroscience research suffer from low power, and especially for published fMRI experiments. Not only does low power decrease the chance of detecting a true effect, it also reduces the chance that a statistically significant result indicates a true effect (Ioannidis, 2005). Put another way, findings with the least power will be the least reproducible, and thus a (prospective) power analysis is a critical component of any paper. In this work we present a simple way to characterize the spatial signal in a fMRI study with just two parameters, and a direct way to estimate these two parameters based on an existing study. Specifically, using just (1) the proportion of the brain activated and (2) the average effect size in activated brain regions, we can produce closed form power calculations for given sample size, brain volume and smoothness. This procedure allows one to minimize the cost of an fMRI experiment, while preserving a predefined statistical power. The method is evaluated and illustrated using simulations and real neuroimaging data from the Human Connectome Project. The procedures presented in this paper are made publicly available in an online web-based toolbox available at www.neuropowertools.org.

## 2 Introduction

In a scientific study, one typically aims for a statistical power of 80%, implying that a true effect in the population is detected with a 80% chance. Power computations allow researchers to compute the minimal number of subjects to obtain the aimed statistical power. As such, power calculations avoid spending time and money on studies that are futile, and also prevent wasting time and money adding extra subjects, when sufficient power was already available.

Prospective power analyses for fMRI experiments have two distinct uses. One is the the optimisation of the experimental design to ensure maximal statistical power for a given scanning duration and various constraints of behavioural paradigms. Methods have been developed to find the optimal number and arrangement of stimuli over the duration of the experiment for each subject (Wager and Nichols, 2003; Friston et al., 1999; Smith et al., 2007). The second use is to find the necessary number of subjects (Desmond and Glover, 2002; Mumford and Nichols, 2008).

While it is straightforward to compute power for a single, univariate response, determining the power of an fMRI study is a formidable task. An array of parameters must be specified such as the within-and between-subject variance, the first and second level design, the temporal autocorrelation and the size of the hypothesized effect, all of which may vary voxel-by-voxel. Many of these parameters may be estimated based on a pilot study, independent of the study to be performed. The most difficult parameter to specify is the region-of-interest (ROI), the specific location where activations are expected.

The earliest attempt to visualise power for neuroimaging was done by Van Horn et al. (1998), where the noncentral F-distribution was used to compute voxelwise power applied to PET data. A first effort to compute sample sizes for neuroimaging for block designs was a procedure by Desmond and Glover (2002), that included within-and between-subject variability and the mean effect. The model was extended by Mumford and Nichols (2008), where multiple designs and temporal autocorrelation were taken into account. The procedure also takes the multiple testing problem into account by averaging over ROI analyses. A more elaborate implementation by Hayasaka et al. (2007) also considered the multiple testing problem by using the non-central random field theory to control the family-wise error rate. In this work we present a simple way to characterize the spatial signal in a fMRI study, and a direct way to estimate power based on an existing pilot study. Specifically, using (1) the volume of the brain activated and (2) the average effect size in activated brain regions, we can directly calculate power for given sample size, brain volume and smoothness. With such a basic formulation, we hope this will make power analyses prevalent, making better use of scarce research funding and better communicating the potential reproducibility of a study.

The present method is an extension of the procedure presented in Durnez et al. (2014) based on peak statistics. Peaks, local maximas in the statistic image, are particularly tractable as they are independent and have reliable random field theory results for their uncorrected and Familywise Error (FWE) corrected *p-*values (Durnez et al., 2014). In contrast, individual voxel values have complex dependency, and clusters have unreliable RFT *p-*values (Woo et al., 2014; Hayasaka, 2003; Durnez et al., 2014; Silver et al., 2011). In our procedure we use a statistic image from a pilot study, and use peaks above a threshold *u* to fit a mixture model, where (1-*π*_1_) of the peaks follow a known null distribution, and the remainder follow a Gaussian with unknown mean and variance. Once the alternative distribution *H*_*a*_ is estimated, the distribution can be transformed to account for a different sample size. As such, not only can the posthoc power of the pilot study be estimated, but also power for a new study with the same experiment and a different sample size, allowing general sample size calculations.

In the remainder of this paper, we present our procedure and evaluate it based on simulations that explore different fMRI characteristics, such as spatial extent of the signal and signal intensity. Next, we present an evaluation using 180 unrelated subjects from the Human Connectome Project (HCP) (Van Essen et al., 2012). These HCP data are used for a number of reasons. First, these data are very high quality, resulting in a very high power when including all subjects and thus offer a high level of certainty about the location of the effect. Second, with 180 subjects, we can use subsamples of the data to create many smaller fMRI studies. The sampled results can then be compared to the results of the full dataset. The added value of the HCP data is that it possesses various unknown noise sources in fMRI that would be impossible to simulate. Third, we demonstrate the procedure on a typical example of an fMRI experiment using an fMRI dataset (Seurinck et al., 2011). Finally, we conclude with a discussion on the topic and the implementation of the procedure in a toolbox.

## 3 Methods

### 3.1 Measures of power

First we consider a single one-sided univariate test: The null hypothesis *H*_0_ is rejected in favor of the alternative hypothesis *H*_*a*_ only when the test statistic *Z* exceeds significance threshold *z*_*α*_; *z*_*α*_ is chosen to control the type I error rate, *α* = *P*(*Z* ≥ *z*_*α*_*|\H*_0_), the power of this test is defined as *P*(*Z* ≥ *z*_*α*_*|\H*_*a*_), which of course requires the distribution of *Z* under *H*_*a*_.

In a multiplicity context like voxelwise testing where many tests are performed simultaneously several definitions for power exist (Dudoit et al., 2003). We will focus on average and familywise power. For voxelwise inference, let 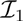 denote the set of indices for voxels that are truly activated. The average power is simply the arithmatic mean of power over non-null voxels:

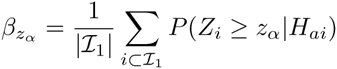

where 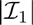 is the number of truly activated voxels, and *H*_*ai*_ is the alternative hypothesis at voxel *i.* Assuming a homogeneous signal, average power has the traditional interpretation of power: the probability of a true positive at one voxel. The familywise error rate (FWER) is the probability of at least one type I error. Its counterpart, familywise power, is defined as *P*(*Z*_*i*_ ≥ *z*_*α*_ for some 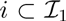), the probability of at least one true positive.

We can similarly define average and familywise power for peakwise tests similar to defining clusters: the identification of local maxima depends on a neighborhood. The SPM^1^ software (RRID:nif-0000-00305) uses 18-order neighborhood to define maxima, while FSL^2^ (RRID:birnlex_2067) uses 26-order neighborhood. While peaks can be defined independently of an excursion or screening threshold, it can be sensible to exclude the lowest peaks with such a threshold *u*. This excludes the peaks corresponding to very low SNR, and also is required if parametric distributional results are to be used. Let 𝒥 comprise the indices of all local maxima above *u*. Denote the test statistic for peak *j* as 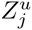. Let 𝒥_1_ ⊂ 𝒥 denote the set of indices for peaks above *u* corresponding to a voxel containing true signal, while 𝒥_0_ ⊂ 𝒥 denotes the set of indices for peaks above *u* corresponding to a voxel containing no true signal. Average power is then defined as

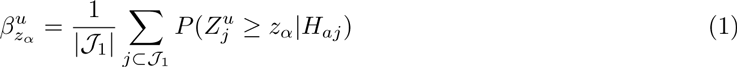

while familywise power is defined as *P*(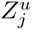 ≥ *z*_*α*_ for some *j* ⊂ 𝒥_1_). In what remains, we focus on average power. First, it is the most intuitive measure of power as it reflects the average probability of a rejection within the set of false null hypotheses. Second, familywise power may not be a satisfactory measure as a high probability of at least one rejection within the set of truly activated peaks may still be accompanied by a large proportion of false negatives.

### 3.2 Estimation Procedure

To calculate average peakwise power, it is essential to know the distribution of peak heights for truly active peaks *Z*^u^ for peak heights above screening threshold *u* (distribution under *H*_a_). Under the null, non-active peaks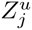 with *j* ∊ 𝒥_0_, have a simple distribution, approximately following an exponential distribution with mean *u* + *1/u* for screening threshold *u* (Worsley, 2007):

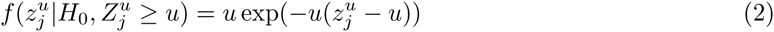

For *H*_*a*_, while a shifted exponential might work well for small signals, we found a truncated normal distribution (truncated at screening threshold *u)* was better to describe the distribution of active peaks 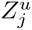 with *j* ∊ 𝒥_1_:

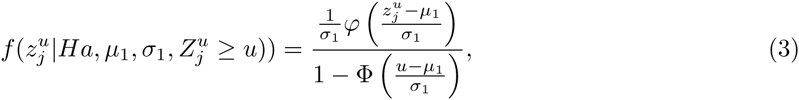

where *ϕ* and Φ are the density and cumulative distribution function of the standard normal distribution, respectively.

The total distribution of peak heights 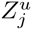 (*j* ∊ 𝒥) can be written as the following mixture distribution:

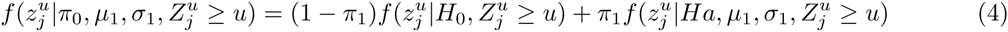

where *π*_1_ is the proportion of true positive peaks among all peaks above *u*.

Instead of estimating all parameters at once, we take a two-stage approach. We first estimate *π*_1_, and then conditional on 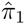 estimate *μ*_1_ and σ_1_. We found this 2 stage approach more stable than estimating all 3 parameters jointly.

There are a variety of estimation methods for *π*_1_ that have been proposed (Benjamini and Hochberg, 2000; Storey and Tibshirani, 2003; Storey, 2002; Pounds and Morris, 2003; Pounds and Cheng, 2004) all based on the observed distribution of the *p*-values. A comparison of the estimators for peak inference in fMRI data analysis is discussed in Durnez et al. (2014), and here we use the preferred method from that work, the estimator of Pounds and Cheng (2004).

Once 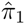 is obtained for a dataset with sample size *n*, we estimate the remaining parameters, *μ*_1_ and σ_1_ in Equation 4 using maximum likelihood on the same data. Let 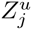 denote an active peak (*j* ∊ 𝒥_1_) with 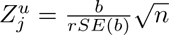, where *b* is the average experimental effect and where *rSE*(*b*) = *SE*(*b*)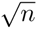. The relative standard error attempts to remove sample size dependence. For a one-sample T-test *rSE*(*b*) equals σ, but this generalises to other types of models and test statistics (see Appendix A). The expected value of the peak height under the alternative before truncation is 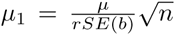 where *μ* = *E*(*b*) is the (non-null) mean in effect units. We define *δ* = *μ/rSE*(*b*) to be the unitless effect size; in the one-sample case, this is exactly Cohen’s d, *δ* = *μ/σ*.

For a new sample of size *n\**, we model the distribution of truly activated peaks before truncation as 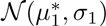 with 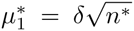. Note that we assume that the variance of the distribution 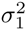 remains constant for different sample sizes.

For a given peak statistical threshold *z*_*α*_, the average peakwise power with truncation at screening threshold *u* is P(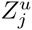 ≥ *z*_*α*_), computed as

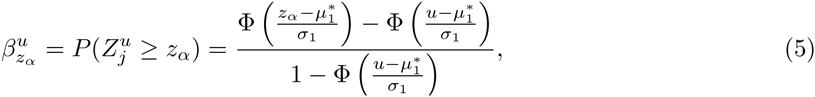

the complementary cumulative distribution function of the alternative distribution from Equation 3. The statistical threshold *z*_*α*_ can either be uncorrected or corrected for multiple testing.

### 3.3 Computing the statistical threshold *z*_*α*_

We evaluate several strategies to select the statistical threshold *z*_*α*_:

1. UN: Level *α* = 0.05, uncorrected for multiple testing.
2. FDR: Corrected to control the false discovery rate at level 0.05 with the method of Benjamini and Hochberg (1995).
3. FWER-BF: Corrected to control the familywise error rate at level 0.05 using a Bonferroni correction.
4. FWER-RFT: Corrected to control the familywise error rate at level 0.05 using a random field theory (RFT) correction (Friston et al., 2007).

The null distribtuion of peaks (Equation 2) is used to compute uncorrected *p-*values for each peak, as well as to compute the thresholds for all of the above methods. We stress that we are not condoning the use of uncorrected thresholds (Method 1), but we need to verify the accuracy of our method in these basic settings. The FDR method (2) is adaptive and produces data dependent thresholds. Our predicted FDR threshold for a new study with *n\** subjects is simply the exact FDR threshold from the pilot study. The rationale behind this is that, given that the actual study is typically larger than the pilot, and that average power will increase with sample size, we expect that the FDR cutoff will decrease. Therefore, using the pilot FDR threshold should in principle result in a conservative estimate of the actually realized FDR threshold, though of course if the pilot is negative (no FDR significance) no power estimation based on FDR can be made. Both methods 3 and 4 assume that the search volume is the same in the pilot and new study, and the RFT method (4) assumes the smoothness is the same.

### 3.4 Simulations

Evaluations are done with 500 simulated datasets. For each realization, for a given sample size *n* we generate *n* statistical parametric maps, summarising evidence for activation. These represent the average effect of the design in a first-level analysis for each subject, i.e. a three-dimensional map of subject-specific b-values. We simulate *n* 64 × 64 × 64 volumes filled with independent standard Gaussian noise, setting the voxel size to 3 × 3 × 3 mm. Images are smoothed with a 3D Gaussian kernel with a full-width at half maximum (FWHM) of 8 mm. After smoothing, images are rescaled to restore unit variance. In each subject-specific b-map, we add true activation in 4 ball-shaped volumes that span either 2, 4, 6 or 8% of the total brain volume. Within these activation blocks, the effect size takes a value of 0.5, 1, 1.5 or 2 units for all *n* subjects. In a next step, a *z* image is obtained by performing an ordinary least squares group analysis with all subject-specific b-maps using FSL’s flame. In this *z* image, peaks are defined and only peaks above screening threshold *u* are considered; we use *u* = 2.3, the standard value in the FSL software package. The null distribution from Equation 2 thus gives peak *p-*values:

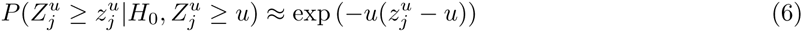

We use these simulations to study the performance of the procedure described in section 3.1. Specifically we first simulate pilot data from 15 subjects. In these data, we estimate 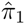 using the procedure presented by Pounds and Morris (2003) and then the alternative truncated normal distribution 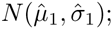 with 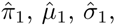, we are able to predict power of future studies in function of sample size *n\**. To estimate the truncated normal distribution using maximum likelihood, we use the Limited-memory Broyden-Fletcher-Goldfarb-Shanno with box constraints (L-BFGS-B) algorithm (Byrd et al., 1995). The expected peak height under the null is *t* = *u* + *1/u* = 2.74. As the expected value of the alternative distribution should exceed this value, we set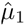 = 2.74 as a lower limit in the optimization algorithm. The standard deviation under the null is *1/u* = 0.43 and we expect the variance under the alternative to be no less than that of the null, so we take 0.1 as a conservative lower limit for 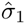. We iterate the optimization over 20 randomly chosen starting values to avoid local maxima in the maximal likelihood estimation. We draw starting values for 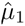 from a range between 2.74 and 10 and for 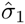 from a range between 0.1 and 10.

For each pilot data set, we predict power for a new study with *n* = 15,…, 35 using equation 5. The true underlying *π*_1_ is obtained by calculating the percentage of peaks that are located in activated areas. The screening threshold has the goal to eliminate a large portion of null peaks. As such *π*_1_ will be slightly higher than the voxelwise proportion of activation in the brain (i.e. 2,4,6 or 8 %). We also calculate the average peak effect size in truly active areas, *E*(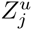),*j* ∊ 𝒥_1_, in the simulated data. We compare this not to 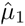, but a shifted version that accounts for the screening threshold *u*, 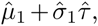 where

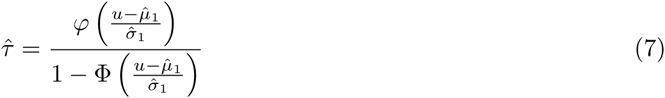

To validate the power predictions for the final experimental study, we simulate study data for *n* = 15,…, 35 as described above. We apply the significance thresholds described in section 3.2 to the revealed peaks. The predicted power 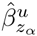 as calculated in Equation 5 is compared to the empiricially derived peakwise average power 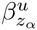 in the simulated images for which the underlying truth is known:

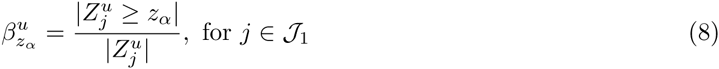

Hence, 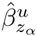 for a sample size *n\** is estimated in each of 500 simulated pilot studies of sample size *n* and the average over these simulations is compared to the average of the actual power in 500 simulated studies of sample size *n\**.

### 3.5 HCP data

We use data from the Human Connectome Project (HCP) (Van Essen et al., 2012) to evaluate our method with complex signal and noise structure. We used results from analysed task fMRI data with 5mm smoothing, ‘level 2’ models (where data from different runs on the same task are combined). For each of 180 unrelated subjects we have 47 unique contrasts.

We use a working assumption that any group analysis of 100 or more participants gives high powered results and can reflect population results.

The procedure used to analyze these data is shown in Figure 1 and is described below. We start with 180 subject-specific b-maps. From this, we take a subsample of 15 subjects that will serve as the held-in pilot data. On these data we perform a Ordinary Least Squares (OLS) group analysis using FSL’s flame and define peaks based on the resulting *z* image. We apply a screening threshold *u* = 2.3, the default value in the FSL software. We derive peaks and compute peak *p*-values (Equation 6) and apply our estimation procedure. As in the simulations, we transform the mean of the alternative distribution to the truncated expected effect size as 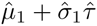 (see Equation 7).

**Figure 1:**
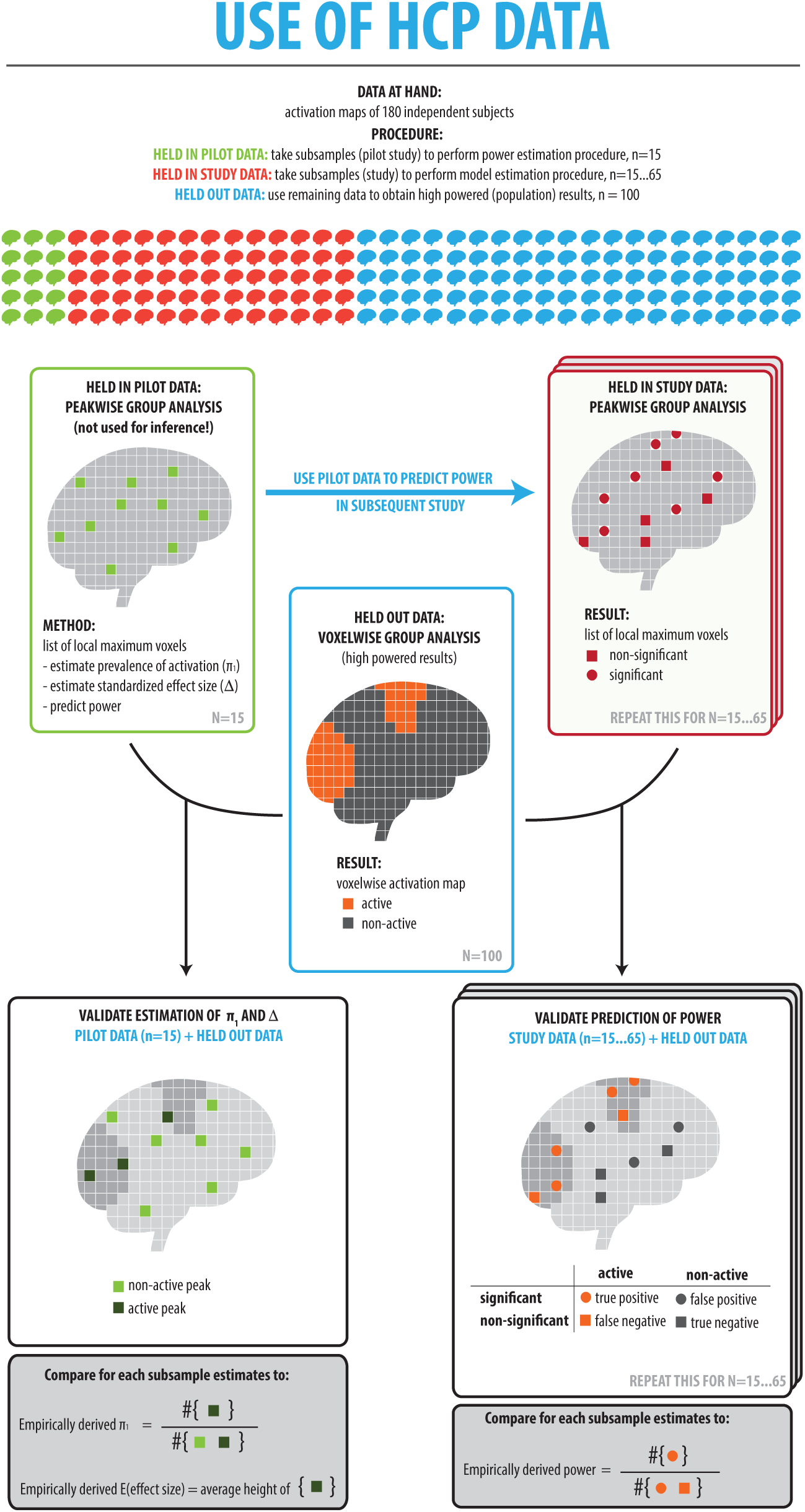
Overview of the procedure used to evaluate power calculations on the HCP-data

Next, we take a subsample of *n* subjects, *n* = 15,11,…, 65. These data represent the experimental data for which the pilot analysis has served. We refer to these data as the held-in study data. With the same procedure as with the pilot data, we compute peak *p*-values and we perform a peakwise analysis with the thresholding procedures described in section 3.2.

On the remaining 100 subjects (the held-out data) we also perform a one-sample T-test, but threshold each *t* image voxelwise at 5% FWER using Random Field Theory (Worsley et al., 1992). With *n*_Ref_ = 100, we will use the set of FWER-significant voxels as working set of **empirically active** voxels 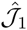, the remaining voxels denoted 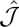_0_.

To validate the estimation procedure, we combine the held-in pilot data and the held-out data to validate the estimation of 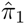 and the effect size as follows: when a peak from the held-in data corresponds to an empirically active voxel in the powerful held-out data, we consider this as an empirically active peak. When a peak corresponds to an empiricially inactive voxel in the held-out data, it is considered empirically inactive. We define **empirically derived** *π*_1_ as the ratio of the number of empirically activated peaks and the total number of peaks (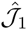/𝒥). The **empirically derived effect size** is the average peak height of all peaks that are located in empirically activated area. Finally, we combine the held-in study data with the held-out data to validate the power estimation: **empirically derived power** is defined as the ratio of the number of significant empirically active peaks for the held-in study data (for a given thresholding procedure) and the total number of empirically activated peaks. The empirically derived power is computed for subsamples with *n* = 15,…, 65 subjects; each power prediction based on *n* subjects is compared to power on these *n* = 15,…, 65 subjects.

This resampling procedure is repeated 500 times for each of the 47 contrasts.

Some tweaking is needed to use these real data for evaluation purposes. While we assume that the analysis of the held-out data is powerful enough to represent results on a population level, these data were analyzed using familywise error rate control. Therefore, the maps with the set of voxels (and consequently peaks) that we call *empirically activated* can be contaminated with 5% false positives. On the other hand, within the voxels (and consequently peaks) that appear *empirically inactive*, there might be false negatives. This poses a problem for our measures of empirically derived *π*_1_, effect size *δ* and power. We have derived a set of corrections for these measures to account for the presence of false positives and false negatives in the held-out data described in Appendix B.

### 3.6 Data example

We apply our estimation procedure to data from a study of 13 subjects on the role of higher order visual areas in imagined visual motion (Seurinck et al., 2011). First level analyses were carried out with a standard General Linear Model approach using FSL’s FEAT function (Jenkinson et al., 2012). The second level analysis is performed in R with OLS, resulting in a 3D T-statistic map, that is transformed to *Z*-values (Hughett, 2007).

For this one Z map we apply our method with *u* = 2, characterizing the signal with 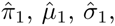 and computing prospective power for different statistical thresholds (uncorrected, FDR-corrected, FWER-corrected with a Bonferroni procedure and FWER-corrected with a Random Field Theory procedure) and sample sizes (*n\** = 10 – 60).

## 4 Results

### 4.1 Simulations

Results for the accuracy of alternative distribution parameters *π*_1_ and *μ*_1_ are shown in Figure 2. We can see that for very small effect sizes, the prevalence of activation *π*_1_ is unbiased, while for medium to large effect sizes *π*_1_ is overestimated. We show in the supplementary materials that this effect can be explained by the mismatch between the modeled beta-distribution and the observed distribution of *p*-values for active peaks.

**Figure 2:**
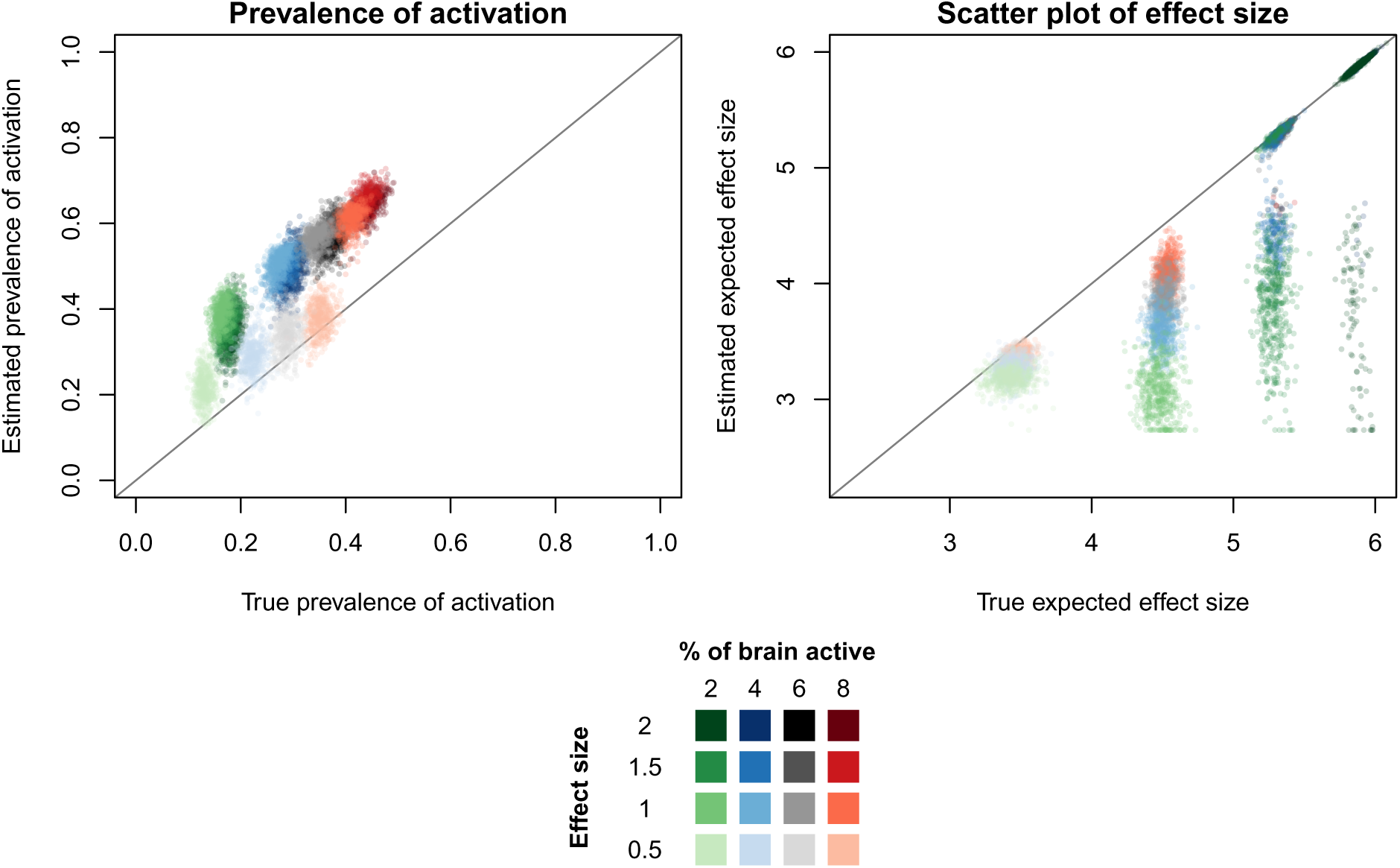
Left: Plot of estimated 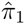 against true *π*_1_> for different sample sizes and different values for *μ*_1_. Each dot represents a different simulation, as such there are 500 dots for each condition. Right: Plot of estimated expected peak height 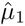 against true expected peak height *μ*_1_ for different effect sizes. The estimations are the result for a pilot dataset with *n* = 15.

The effect size estimator is unbiased for the smallest effect size. The larger the effect size, the better the estimation. When only few activation areas are present, the effect size is underestimated. In this case, the number of activated peaks is very small in relation to the number of null peaks. Therefore, the model shows biased results in separating both distributions. With larger effect sizes and larger activation areas, the model performs very well.

Using these estimates of 7Ti, *π*_1_, *μ*_1_ and σ_1_ for the case of *n* = 15, we then computed power for future studies for peak inference with 5% error rate control for the different inference procedures tested. Figure 3 shows the plot of 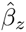 and *β*_*z*_ for *n\** = 15 – 35. With small activated brain regions, the power is largely underestimated. This leads to conservative results. For larger activated regions, the results are unbiased. This result is consistent over different conditions and different thresholding procedures. The procedure is often unable to estimate power for FDR thresholding for smaller effect sizes. This is because - due to the adaptive character of the procedure - in the pilot study often no significant effects are found. Therefore, there is no cutoff, and as such power cannot be estimated in larger sample sizes. The non-adaptive multiple testing procedures show a modest bias only.

**Figure 3:**
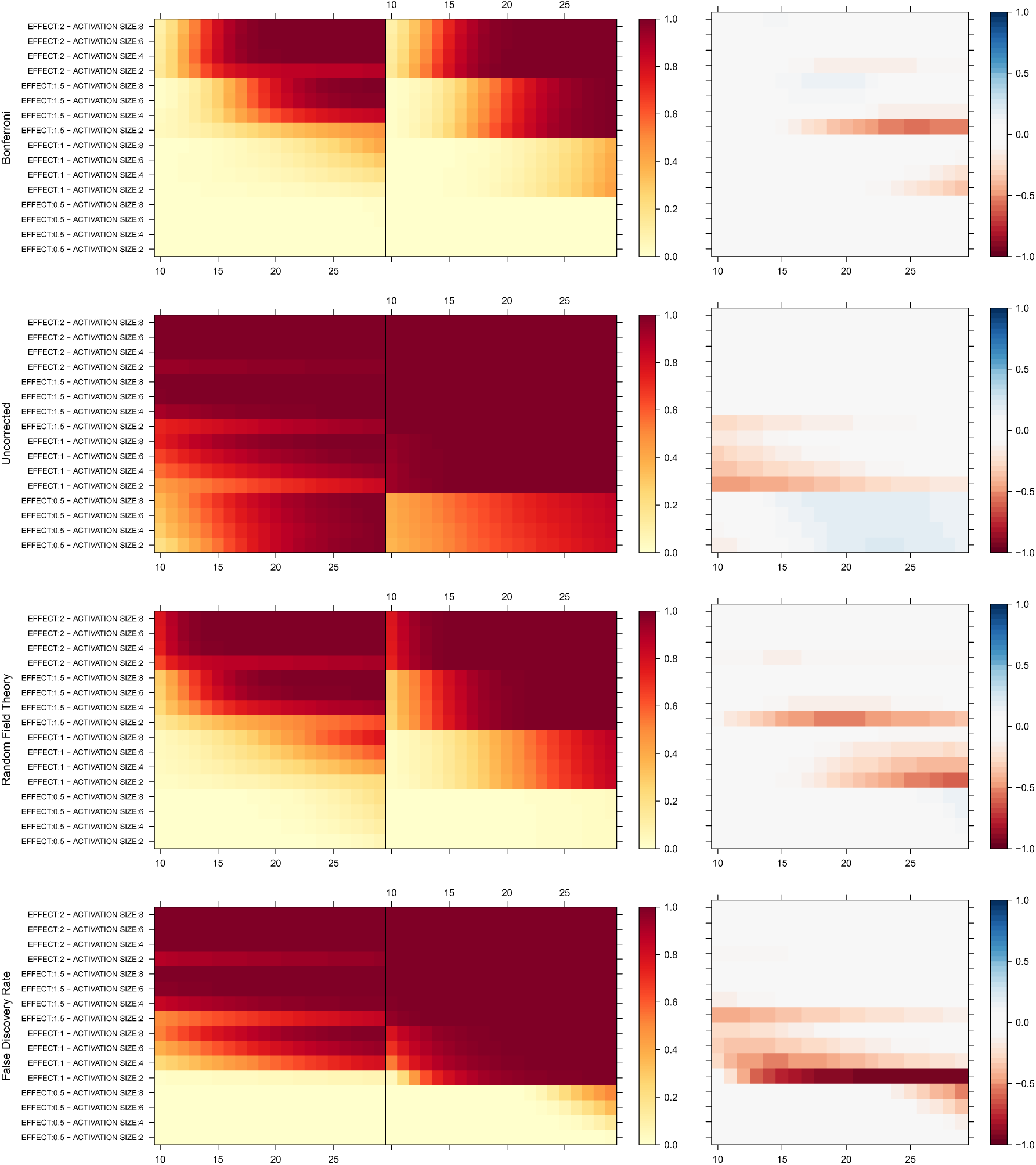
Plots of the peakwise average power with error rate control at 5% for different effect sizes and different amounts of activation. The left column shows the estimated power curves, the middle column shows the true power and the right column shows the bias. Bias is defined as the estimated power minus the true power. The peakwise average power is estimated from a pilot study with 15 subjects.

The goal of the presented method is to perform sample size calculations. Figure 4 shows the performance of these sample size calculations when aiming for 70% power. While the variance is high for conditions with small effect sizes, the average bias of most conditions and multiple comparison procedures is within a range of 5 subjects. In line with the previous results, the model has difficulties estimating power when only a small proportion of the brain is active.

**Figure 4:**
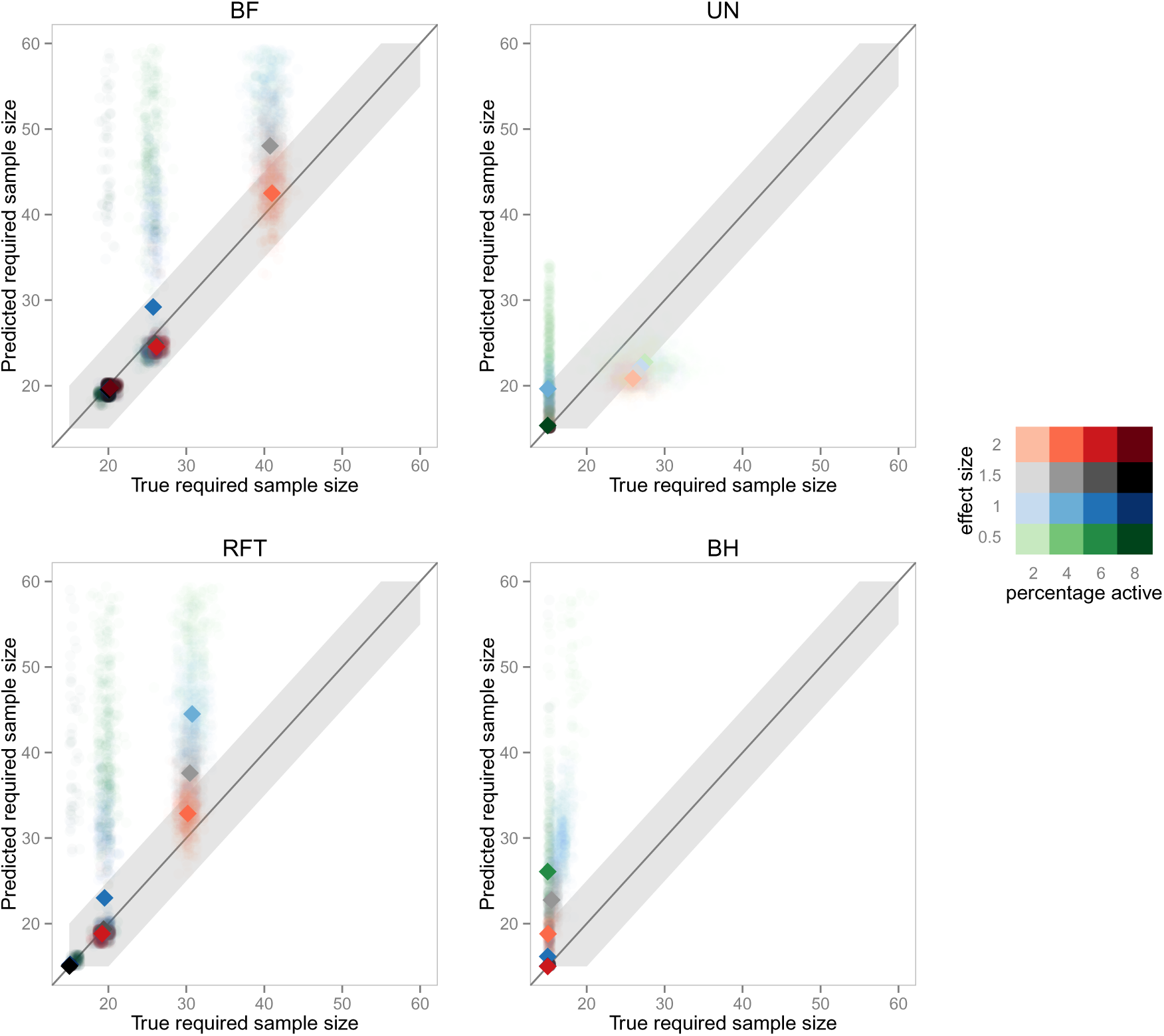
Plots of the predicted and true required sample size when aiming for 70% power. The different plots refer to the different multiple testing procedures. Points inside the grey area identify poins with a maximum bias of 5 subjects. Each semi-transparent dot represents a different simulation, as such there are 500 dots for each condition. The fully colored dots present the average per condition. The estimated sample size results from a pilot study with 15 subjects.

### 4.2 HCP data

Results of the estimation procedure for the prevalence of activation *π*_1_ and the effect size based on a pilot dataset of about 15 subjects are shown in Figure 5. We show that for a range of contrasts and tasks we find slightly overestimated estimates for the prevalence of activation and the effect sizes. The estimations are good with low variance and the effect sizes suffer from modest overestimation. The result of the gambling task contrast, “punish versus reward” is noteworthy. There are very few activations in this contrastoften leading to no active peaks. Only in 19 of the 500 pilot data resamples, there were peaks in the activated area, as such the average true *π*_1_ and 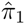 are very close to 0. However, even in this contrast, the effect size is well estimated in the cases where 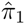 was above 0.

**Figure 5:**
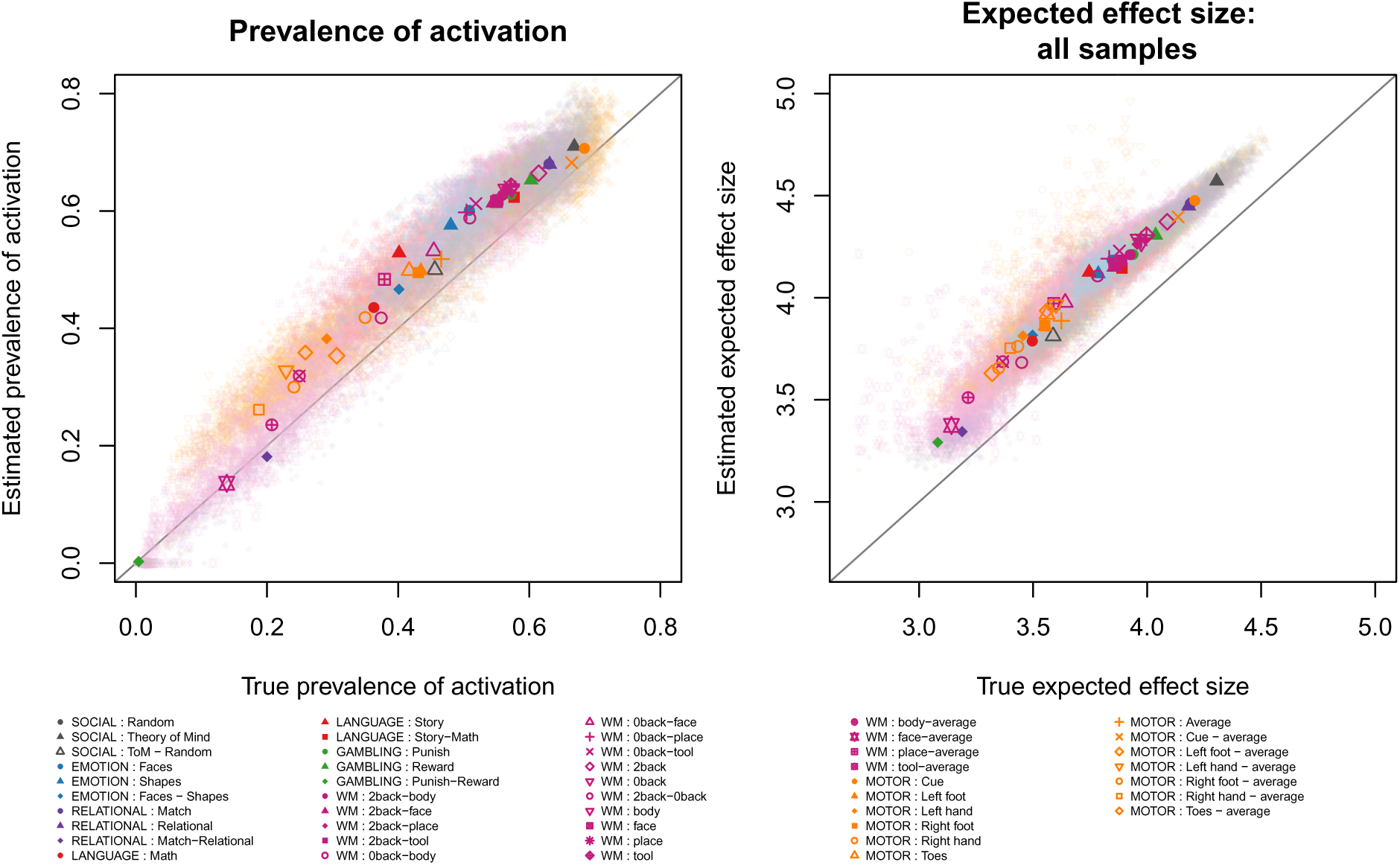
An evaluation of the estimation of *π*_1_ (left) and the effect size (right). Each pastel colored plotting symbol corresponds to one random sub-sample taken from the data, for one particular sample size, experiment and contrast. The average estimation for each contrast is plotted in a darker color.

The power estimates are accurate over a large range of tasks. The estimation and bias on the power estimates for error rate control at 5% is given in Figure 6. In most contrasts, the bias ranges from 10% underestimation (blue) to 10% overestimation (red). In general, the underestimation disappears over a short range of subjects, indicating that a sample size calculation will overestimate the number of subjects needed, but by only a few subjects difference. The underestimation is much larger for contrasts with a smaller effect size. As in the simulations, the adaptive multiple testing procedures show instability due to instances when no significant results are found, leading to underestimation of power. All power estimates are at most conservative.

**Figure 6:**
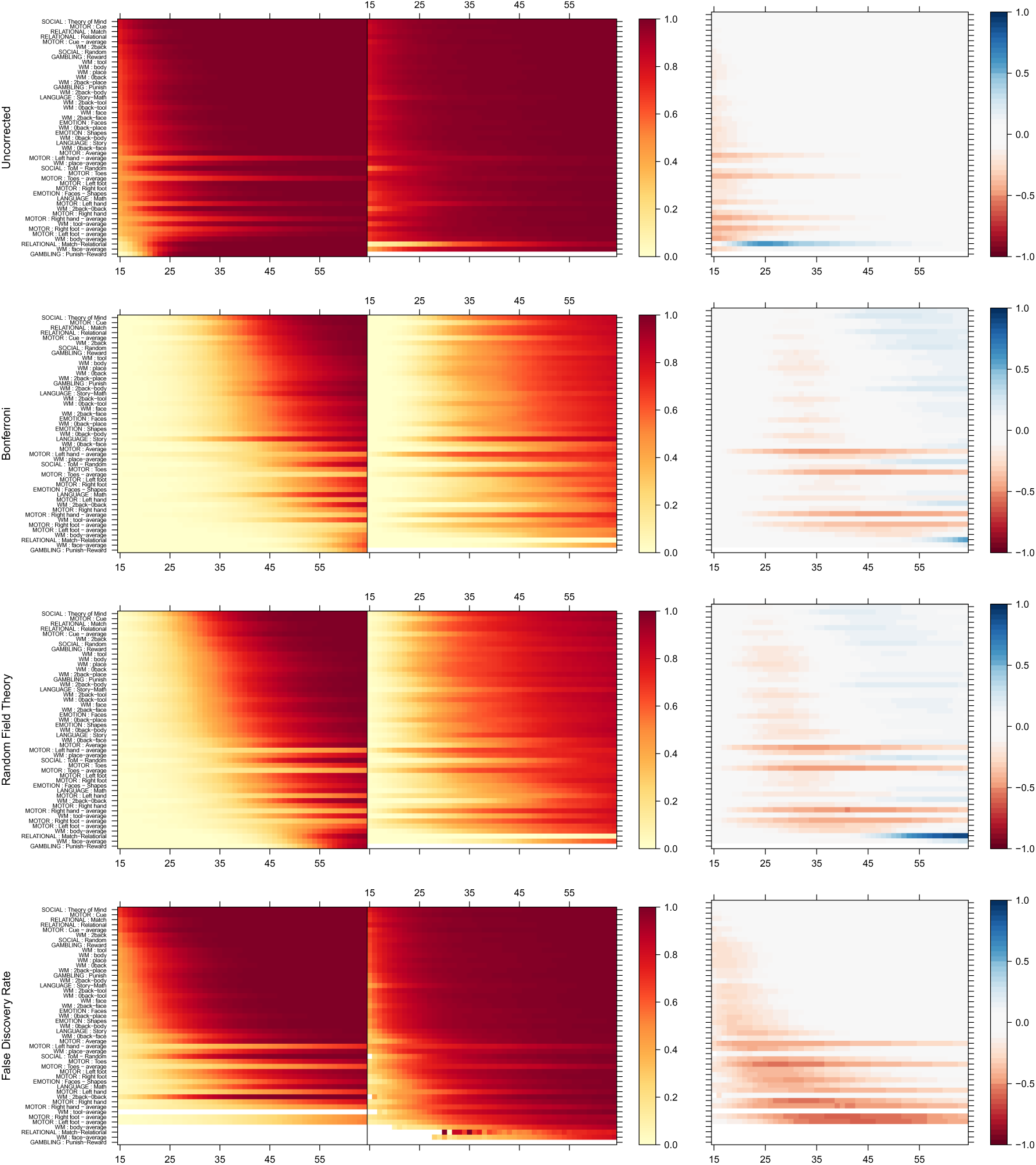
Evaluation of the power estimation over different subjects for all unique HCP-contrasts for thresholding with different error rate corrections at *α* = 0.05 from a pilot study with 15 subjects. The left column shows the estimated power curves, the middle column shows the true power and the right column shows the bias. Bias is defined as the estimated power minus the true power. The contrasts are sorted by their average empirically derived effect size.

Again, the gambling task contrast, “punish versus reward”, is problematic. There are no resample schemes where both the pilot study resamples and the final study resamples result in the detection of active peaks. As there is no activation, there is no finite solution for the average power defined in equation 1. The same problem arises for some contrasts when thresholding with false discovery rate control. As in the simulations, due to the adaptive character of the procedure, sometimes no significant effects are found. This leads to infinite computations for power.

Figure 7 shows the performance of the model when predicting sample sizes. We see that our method predicts very well the sample size for a large range of contrasts. Only the motor contrasts show a larger bias. This can be due to the fact that the motor contrasts are known to be very local contrasts. This corresponds to the results from the simulated condition when the percentage of active brain regions is small.

**Figure 7:**
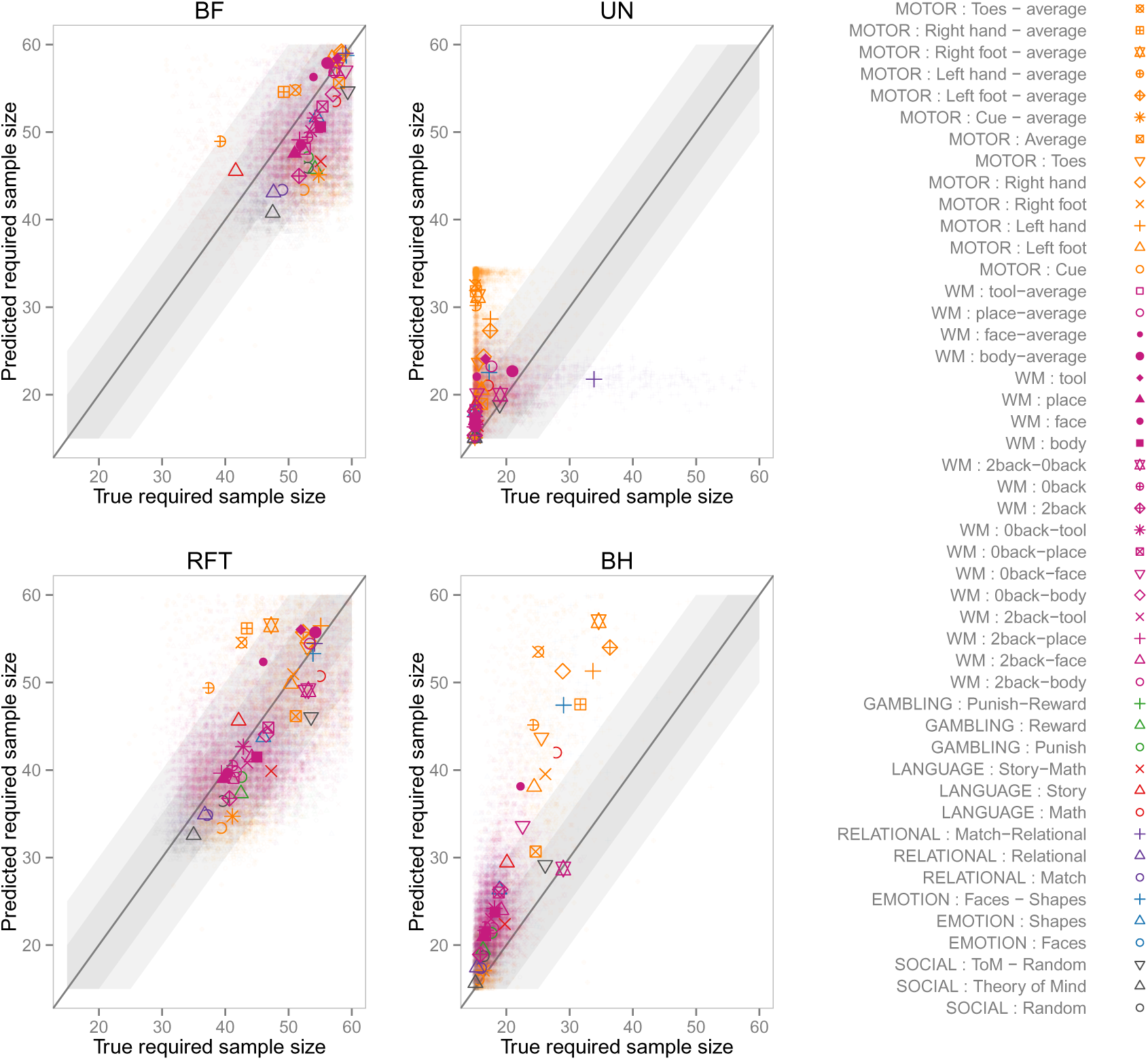
Plots of the predicted and true required sample size when aiming for 70% power. The different plots refer to the different multiple testing procedures. Points inside the light grey area identify poins with a maximum bias of 15 subjects, the darker grey area refers to a maximum bias of 5 subjects. Each semi-transparent dot represents a different subsample, as such there are 500 dots for each condition. The fully colored dots present the average per task. The estimated sample size results from a pilot study with 15 subjects.

### 4.3 Example

Based on these values of *π*_1_, *μ*_1_, *σ*_1_, power predictions for different thresholding methods and a range of *n*\*’s are sho in Figure 8.

**Figure 8:**
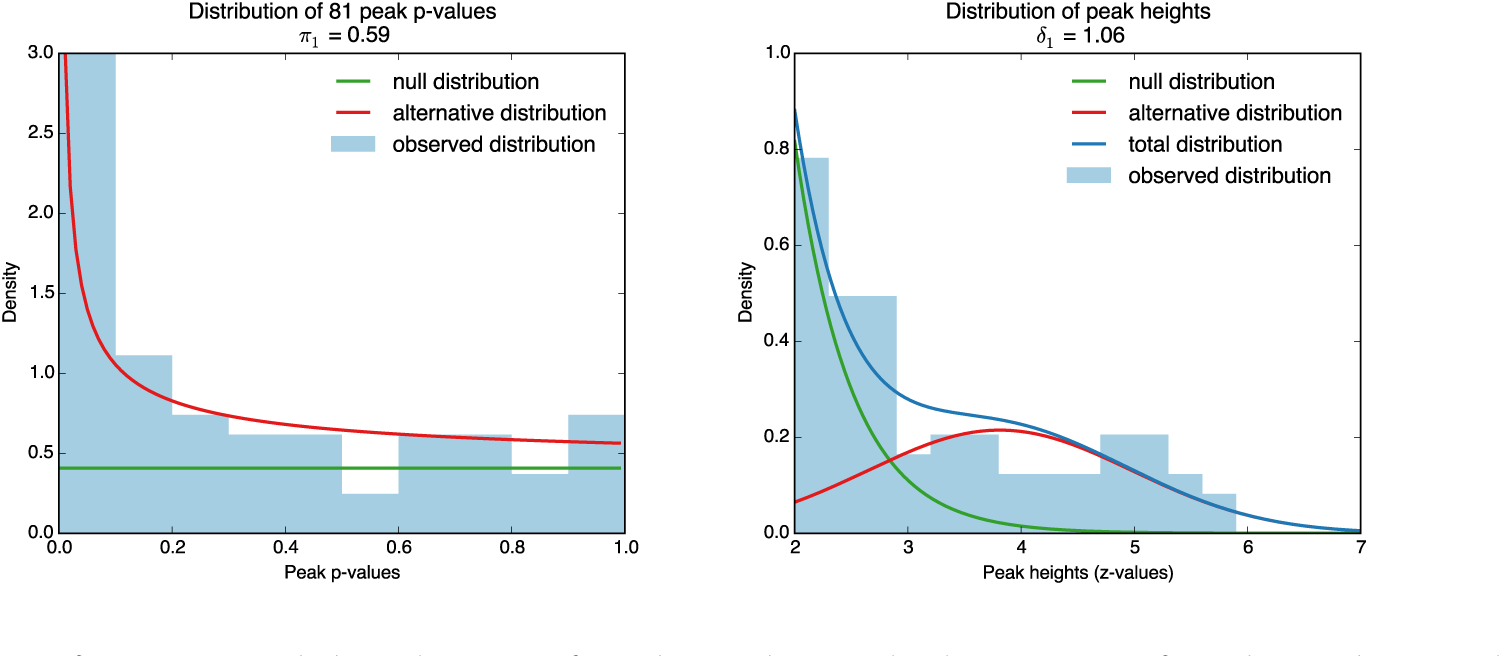
Left: Estimated distribution of peak *p*-values. The histogram of peak *p*-values is shown in light blue, the lines show the estimated part of the histogram stemming from the null distribution (green) and the total distribution (blue). Right: Estimated distribution of peak heights. The histogram of the peak heights is shown in light blue, the lines show the estimated distributions for the null (dark green), the alternative (light green) and the total distribution (blue)

The results of the power estimation procedure can be found in Figure 9. If the aim of the power analysis is to obtain at least 80% average power, then this study would require 19 subjects for uncorrected thresholding at *α* = 0.05. False discovery rate thresholding at *α* = 0.05 requires 27 subjects. For family-wise error rate control at *α* = 0.05, the study would require 41 subjects with Bonferroni thresholding and 42 subjects with Random Field Theory thresholding.

**Figure 9:**
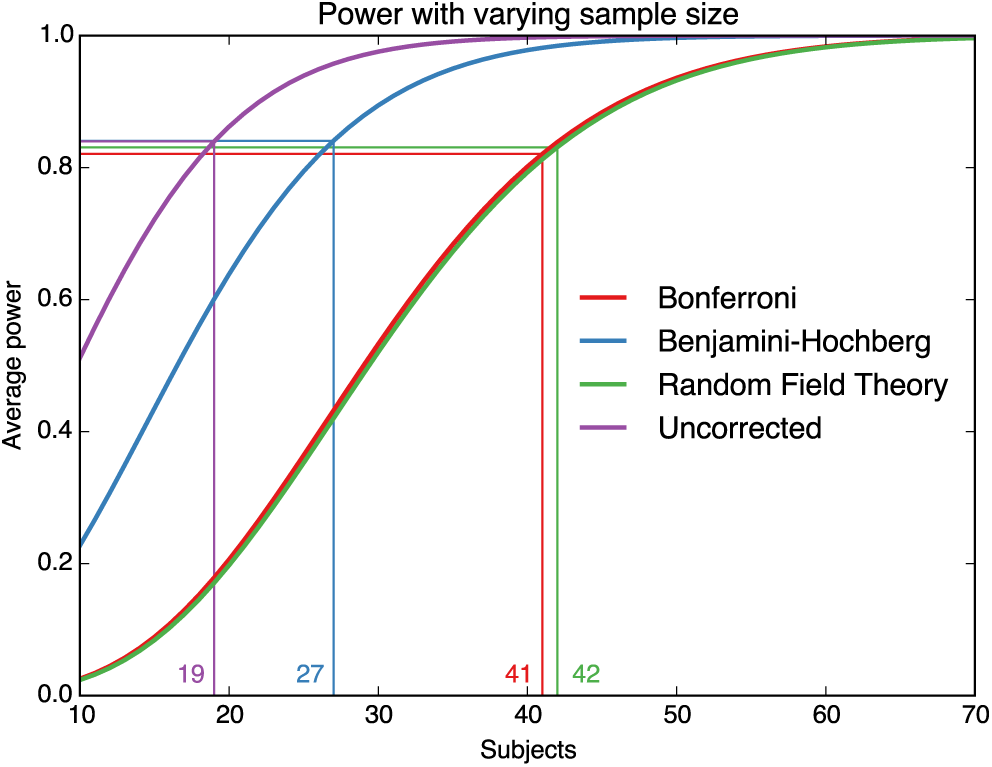
Estimated peakwise average power, *u* = 2 for different multiple testing procedures as a function of sample size. The vertical lines show for each multiple testing procedure the required number of subjects to obtain a power of at least 80%.

## 5 Discussion

In neuroscience, results are often based on fMRI studies that suffer from a lack of power. In order to save costs and effort while preserving sufficient power for detecting important effects, we presented a method to predict power for different sample sizes. While other methods for power calculations in fMRI often require the estimation of many different parameters that are often difficult to estimate (Mumford and Nichols, 2008; Desmond and Glover, 2002), our method is based on only peak values that require Random Field Theory assumptions for the computation of *p*-values.

Our results indicate that the method works well when the size of activated brain regions is reasonably large. When the activated region is small, the presented method underestimates the power largely. This can be seen in the simulations, where biases arise when only 2% of the brain is active but disappear when over 6% is active. In the HCP data, we also see larger biases for the motor tasks, a contrast known for larger effect sizes but very local effects. This indicates further that biases arise when the activated region is small, irrespective of the effect size. Applying the method to small activated brain regions results in atmost conservative results for sample size calculations, i.e. overestimation of required sample size. However, to confidently use the method, it is best to restrict the search region (i.e. apply a ROI mask), an advice that generally applies to fMRI analysis.

Furthermore, the pilot study should be sufficiently high for the power analyses to work. In the supplementary materials, we show the results for a smaller sample size (*n* = 10), which results in larger biases for small effect sizes and small activation foci.

We focus on peaks for several reasons. We have shown in previous work that the assumption of a uniform distribution of the p-values under the null is mainly attained with peak *p*-values, but not with cluster *p*-values (Durnez et al., 2014). As this is an assumption crucial for the procedure presented here, we opt for peak inference rather than cluster inference. Moreover, problems with localisation and stability have been reported with cluster inference (Roels et al., 2014; Woo et al., 2014). However, when a user wants to infer power for cluster inference, this procedure on peaks can be used as a lower bound, as the power of cluster inference should be generally higher than peak inference (Friston et al., 2007). While voxelwise inference was not directly considered, RFT FWER inferences for peaks should give similar results as voxelwise inference, making our power predictions relevant for that domain as well. On the other hand, it is possible to use the method if the prevalence of activation is widespread in the brain. It is up to the researcher to decide whether this shall be the call in the new study.

We have evaluated the procedure using simulated data. The data represent a simplified fMRI experiment but we still vary a number of parameters, like the effect size and the thresholding procedure to ensure that the findings are generalizable to a range of different possible fMRI experiments. In our simulations, we have used a constant effect size of activation over different subjects. We have not applied subject-specific effect sizes, as we believe this would not alter the average effect size, but rather it would inflate the between subjects variance, leading to a smaller normalized effect size. Thus we have considered only varying the average effect size *μ*_1_ and not separately the between subject variance.

This method is only a first step in developing a means to better predict the power of fMRI studies. Many different extensions are possible. One of these possibilities is the development of a testing procedure that would allow to use the pilot data in the final study without harming the false positive rate (see, e.g., similar ideas in genetics Skol et al. 2006). Second, the estimated effect size could incorporate other characteristics besides sample size, like intrasubject variance or scan time (Mumford and Nichols, 2008) These additional parameters would allow the optimization of future studies without the restriction that all characteristics are identical to the pilot study.

Although the evaluation on this method was performed on whole-brain analyses, it is also possible to only apply it to a certain part of the brain, when a region of interest is specified.

We have made the procedure available to the community in a toolbox which is publicly available at http://www.neuropowertools.org, for which the code can be found at https://github.com/neuropower/neuropower. All code used for the validations and example in this paper are available online http://github.com/jokedurnez/neuropower-validation/.

## Acknowledgements

We would like to thank Dr. Deanna Barch and Dr. Greg Burgess for their kind help in harvesting the HCP data and comments. This work was partially supported by the Laura and John Arnold Foundation and the Research Foundation - Flanders (FWO). The computational resources (STEVIN Supercomputer Infrastructure) and services used in this work were kindly provided by Ghent University, the Flemish Supercomputer Center (VSC), the Hercules Foundation and the Flemish Government and department EWI. We would like to thank Stanford University and the Stanford Research Computing Center for providing computational resources and support that have contributed to these research results.

## Appendix

**A. Generalisation from one-sample *T*-test to other models**

In section 3.2 we declare how all *Z* statistics can be written in the form 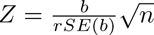 where *rSE*(*b*) = *SE*(*b*)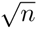. For a one sample *t* test *rSE*(*b*) = σ. Here we explain how this approach can be applied to other types of models.

**A.1 Two-sample *T*-test**

For a two-sample *T*-test, the *Z* statistic is:

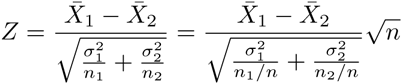

where 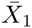 and 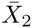 are the sample means of each group, 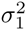 and 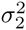 are the respective variances, *n*_1_ and *n*_2_ are the respective sample sizes and *n* = *n*_1_ + *n*_2_ is the total sample size.

Here 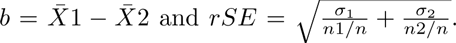 The relative SE can be seen as only depending on the relative sample sizes, and thus the computed power set a total sample size and will be appropriate for a new study that has the same group variances relative sample sizes.

**A.2 Linear regression**

In linear regression, the statistic *Z* for a regressor *X* can be written as:

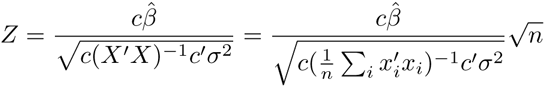

Thus 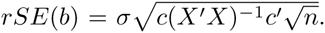 But note that this can also be written as 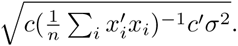 If we regard *x*_*i*_ ~ *F*, then the *p* × *p* term in the denominator can be seen as an uncentered second moment:

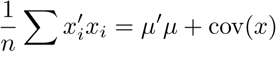

where *μ* = *E*(*x*). Thus the sample size can be set arbitrarily, with the assumption that new observations of *x* can be drawn from *F*.

**B Corrections for the mismatch between true and empirically derived *π*_1_, effect size and power**

We first introduce some notation in table 1:

**Table 1:**
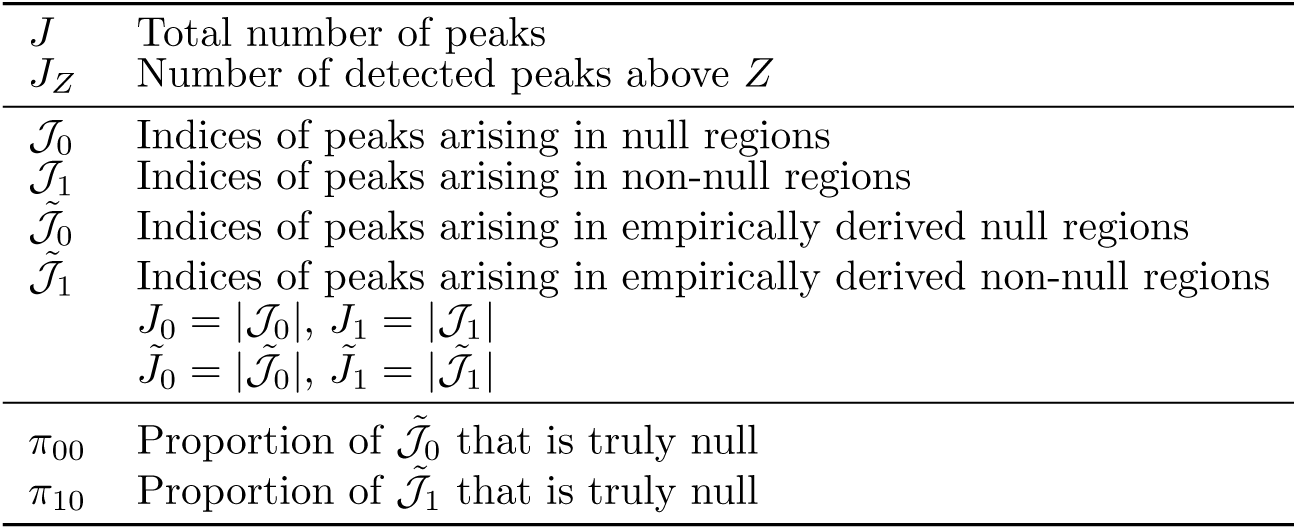
Notation for correction of population level estimators of *π*_0_ and *μ*_1_.

*π*_00_ and *π*_10_ can be estimated using the Beta-Uniform Model by Pounds and Cheng (2004). With these definitions, we can derive the number of peaks that are falsely classified in the FWE-analysis in table 2:

**Table 2:** Classification table of peaks after FWE thresholding.

|  | Empirically derived<br>Null peaks | Empirically derived<br>Active peaks |  |
| --- | --- | --- | --- |
| True null peaks | $\pi_{00}\tilde{J}_0$ | $\pi_{10}\tilde{J}_1$ | $J_0$ |
| True active peaks | $(1 - \pi_{00})\tilde{J}_0$ | $(1 - \pi_{10})\tilde{J}_1$ | $J_1$ |
| | $\tilde{J}_0$ | $\tilde{J}_1$ | |

Thus, an uncontaminated estimate of *π*_0_ can be found as:

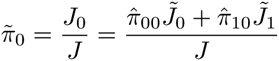

Similarly, bias-corrected versions of *μ*_1_ are possible. First we note

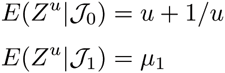

where the conditional expectation indicates the set of peaks under consideration. Now, similar to table 2, we can decompose the expectation over observable sets:

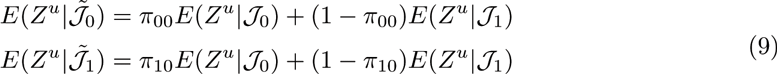

Thus 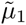 can be estimated:

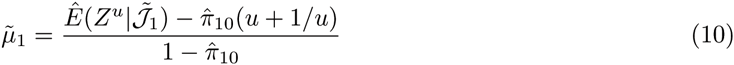

## Footnotes

1 http://www.fil.ion.ucl.ac.uk/spm/

2 www.fmrib.ox.ac.uk/fsl

